# A morphological analysis of activity-dependent myelination and myelin injury in transitional oligodendrocytes

**DOI:** 10.1101/750083

**Authors:** Eszter Toth, Sayed Muhammed Rassul, Martin Berry, Daniel Fulton

## Abstract

Neuronal activity is established as a driver of oligodendrocyte (OL) differentiation and myelination. The concept of activity-dependent myelin plasticity, and its role in cognition and disease, is gaining support. Methods capable of resolving changes in the morphology of individual myelinating OL would advance our understanding of myelin plasticity and injury, thus we adapted a labelling approach involving Semliki Forest Virus (SFV) vectors to resolve and quantify the 3-D structure of OL processes and internodes in cerebellar slice cultures. We first demonstrate the utility of the approach by studying changes in OL morphology after complement-mediated injury. SFV vectors injected into cerebellar white matter labelled transitional OL (_T_OL), whose characteristic mixture of myelinating and non-myelinating processes exhibited significant degeneration after complement injury. The method was also capable of resolving finer changes in morphology related to neuronal activity. Prolonged suppression of neuronal activity, which reduced myelination, increased the number of _T_OL processes, while decreasing both the length of putative internodes, and the proportion of myelinating terminal branches. Overall this approach provides novel information on the morphology of _T_OL, and new opportunities to study the response of OL to conditions that alter circuit function or induce demyelination.

## INTRODUCTION

The speed of information transfer in axons is enhanced by myelination, with processing speed rising as axonal ensheathment increases [1, 2]. These gains in signal transmission arise as a result of alterations in the electrical properties of myelinated nerve fibre, including reduced transmembrane capacitance and increased resistance, which collectively increase the axonal length constant, allowing depolarising current to spread over greater distances before requiring a boost from active depolarising currents. Consequently, conduction velocities in myelinated axons are enhanced significantly over those of unmyelinated axons of a similar caliber once a specific diameter, estimated at 0.2 µm, is exceeded [3]

Myelin formation and neuronal activity are intimately linked. Pharmacological blockade of neuronal activity reduces myelination in numerous *in vitro* and *in vivo* systems [4–6], while increased neuronal activity, induced either optogenetically [7], or *via* sensory input [8], has the opposite effect. This connection between circuit function and myelination, and the eventual contribution myelination makes to functioning of the recipient axon, has led to the concept of myelin plasticity, and an appreciation of the contribution it may make to cognitive function and dysfunction [9–12]. To study myelin plasticity, and evaluate the cellular and molecular mechanisms driving its expression, it is necessary to develop methods that enable a dynamic analysis of internode formation and characteristics under differing levels of axonal activity. Internode thickness and length are particularly relevant to these analyses [13]. Increases in myelin thickness simultaneously reduce the transmembrane capacitance charge, and the leak of cations from the axoplasm towards the extracellelular space, which together increase the spread of depolarizing current along the axon towards the next node. Consequently, the axonal length constant increases allowing depolarizing current to travel further before requiring time and energy consuming boosts from active nodal currents. Intermodal length is therefore free to increase so that enhancements in conduction velocity scale with increasing internode length until they reach a ‘flat maximum’ from which no additional increases can be achieved [14]. Internode thickness and length are therefore important factors in determining axonal conduction velocities.

Although internode thickness is a key parameter controlling conduction velocity, its measurement is unsuited to a dynamic analysis, for example by live-imaging, since it requires the histological analysis of transected axons. Internode length, on the other hand, can be examined in transgenic reporter mice in which fluorescent protein (FP) expression is targeted to oligodendrocyte lineage cells (OL), or in OL labelled by FP expressing viral vectors. In the former context, longitudinal *in vivo* two-photon microscopy has been used to visualise the morphologic responses of adult OL internodes labelled by membrane targeted green FP (mGFP) [15]. Interestingly, environmental enrichments designed to provoke increased neuronal activity, induced an increase in newly formed OL without affecting internode dynamics [15]. Indeed, the internodes of mature OL maintained a remarkable stability even when monitored over 50d. This stability in OL morphology is not universal however since sensory deprivation reduces OL process length and internode number in the cortex of newly weaned mice (21 days) [8]. OL therefore seem to exhibit a greater degree of morphological plasticity at earlier stages of CNS development. Indeed, data from neonatal cortical slice cultures reveal a reduction in internode length following GABA-A receptor blockade [16]. It is important to note that the majority of studies in this area have focused on cortical gray matter OL [7, 8, 15], and that *in vivo* imaging of deeper lying white matter structures represents a significant challenge to current imaging methodologies [17]. For these reasons, the question of how neuronal activity alters OL and internode morphology, and how these effects vary across developmental ages and anatomical regions, for example grey and white matter, remains an open question.

In this paper, we report the morphological analysis of neonatal white matter OL within a system amenable to prolonged pharmacological manipulations capable of either modulating neuronal activity or inducing OL myelin injury. We adapted an approach involving the gliotropic subtype of the Semliki Forest Virus (SFVA7(74) (SFV) [18], previously used for labelling OL in cultured hippocampal slices [19], to label OL in the white matter of neonatal cerebellar slice cultures. Injection of SFV vectors encoding membrane targeted FPs into cerebellar white matter provided an efficient and rapid labelling of myelinating OL, the brightness and extent of which allowed the complete resolution of complex OL process arbors. OL image stacks were traced and the resulting 3-D reconstructions analysed to provide quantitative data on OL process arbor morphology. Using this approach, we analyzed OL process length and branching, internode number and length, and the number and relative proportions of myelinating and non-myelinating branches. The results provide new information on the morphology and activity-dependent development of transitional OL (_T_OL). _T_OL are an intermediate stage of maturation linking premyelinating OL, in which all process branches (PB) are non-myelinating (_NM_PB) [20], and mature myelinating OL (_M_OL), in which it is expected that all PB are myelinating (_M_PB). In this work, _T_OL are defined as cells possessing a mixture of _NM_PB and _M_PB [19, 20], and the term myelinating OL is used generically to include both _T_OL and _M_OL.

We have confirmed that this methodology resolves changes in _T_OL morphology in slice cultures subjected to a complement-mediated injury. 3-D reconstructions obtained after complement injury revealed clear evidence of damage, including the fragmentation and loss of _T_OL processes and internodes, consistent with myelin damage observed after MBP staining. To analyse changes in morphology associated with neuronal activity, slices were subjected to a chronic treatment with tetrodotoxin (TTX). Immunohistochemical analysis showed that TTX treatment reduced myelination confirming that neuronal activity influenced myelination in cerebellar slice cultures [5]. This reduction in myelination was accompanied by changes in _T_OL morphology that included an increase in the number of process arbors and a reduction in the length of putative internodes supported by these processes. Interestingly, TTX increased the total number of PB, and the proportion of PB that were _NM_PB, implying that basal neuronal activity directs the differentiation of _T_OL processes towards the production and elongation of internodes. Overall, this work demonstrates the suitability of brain slice cultures for the study of OL morphological plasticity and injury, and provides the foundation for work to track the dynamic response of OL processes arbors and internodes to changing physiological conditions, or pathological insults, in deep lying white-matter regions that are currently beyond the range of *in vivo* imaging methods.

## RESULTS

### SFV vectors label myelinating OL in the white matter of neonatal cerebellar slice cultures

In our previous work, we applied SFV vectors encoding farnesylated FP (eGFP-f / mCherry-f) directly to the surface of cerebellar slice cultures to label glia [5]. Owing to the limited diffusion of viral particles through the tissue slice [21], this method predominantly labelled glia residing close to the upper surfaces of the slice, including NG2^+^ OPC and astrocytes, but very few cells with a morphology typical of myelinating OL. In the present work, we attempted to label myelinating OL in deeper regions of the slice by microinjecting SFV vectors into the white matter tracks of the folia (Fig. 1Ai) where myelinated Purkinje cell axons reside (see Supplementary Materials and Methods online). In agreement with previous studies in hippocampal slice cultures [19, 22], these injections labelled glia with multiple parallel-aligned processes typical of myelinating OL (Fig. 1Aii). Immunostaining with antibodies to MBP (Fig. 1B) and MOG (Fig. 1C) revealed positive localization of these myelin proteins to SFV labelled OL (Fig. 1B; 2C), while staining of axons with anti-NF200 revealed close alignments of axons and the processes of SFV labelled OL (Fig. 1D). In addition SFV vectors illuminated the myelinating morphology of OL expressing CNPase-GFP transgenes [23] (see Supplementary Fig. S1 online). Taken together these data confirm that SFV labelling identified myelinating OL in the cerebellar white matter.

**Figure 1.**
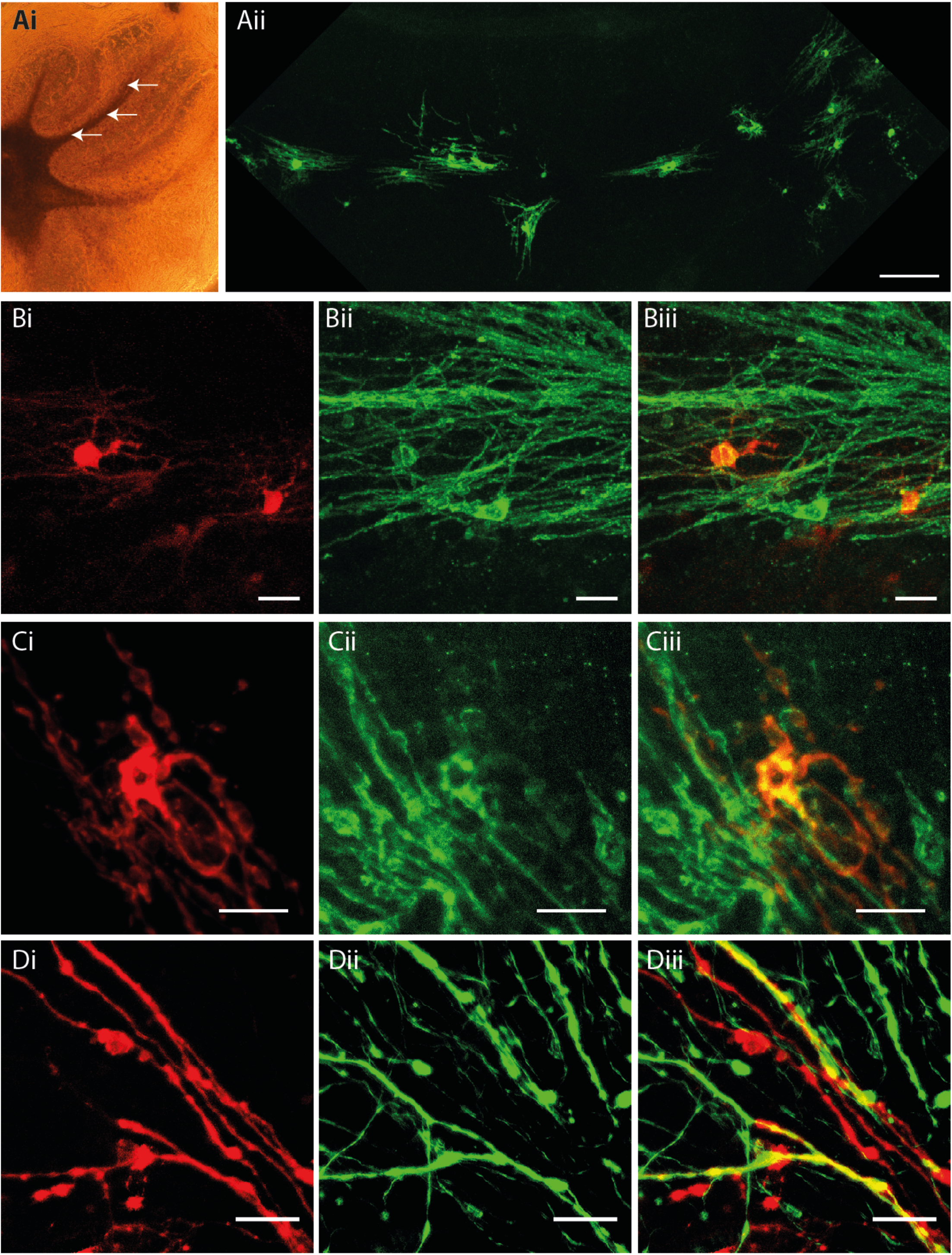
SFV transduces myelinating OL in cerebellar slice cultures. Ai. Phase contrast image of a cerebellar slice culture illustrating white matter tracks and typical sites for SFV injections (arrows). Aii. OL with a myelinating morphology are labelled throughout the entire white matter track of one folium 24 hours after injection of SFV encoding eGFP-f. Magnification 10x, Scale bar 100µm. B. SFV labelled OL express MBP. Bi. Cytoplasmic mCherry in two white matter cells with typical myelinating OL morphology. Bii. MBP immunofluorescence in the field shown in Bi. Biii. Merged image indicating localization of anti-MBP (green) to the processes of the mCherry^+^ (red) cells depicted in Bi. Magnification 20x, Scale bars in B panels 20µm. C. SFV labelled cells localize MOG. Ci. mCherry-f labelled white matter glial cell with myelinating morphology. Cii. MOG immunofluorescence in the field shown in Ci. Ciii. Combined mCherry and MOG signals indicate colocalization of anti-MOG (green) and mCherry (red) signals. Magnification 40x, Scale bars in C panels 20µm. D. Axonal interactions of SFV labelled OL. Di. High power (40x) image of processes (red) extending from an OL transduced with SFV-mCherry-f. Dii. Axons (green) from the field depicted in Di labelled with anti-NF200. Diii. Merge image indicating localisation of NF200 on SFV labelled processes (yellow). Scale bars in D panels 20 µm.

**Figure 2.**
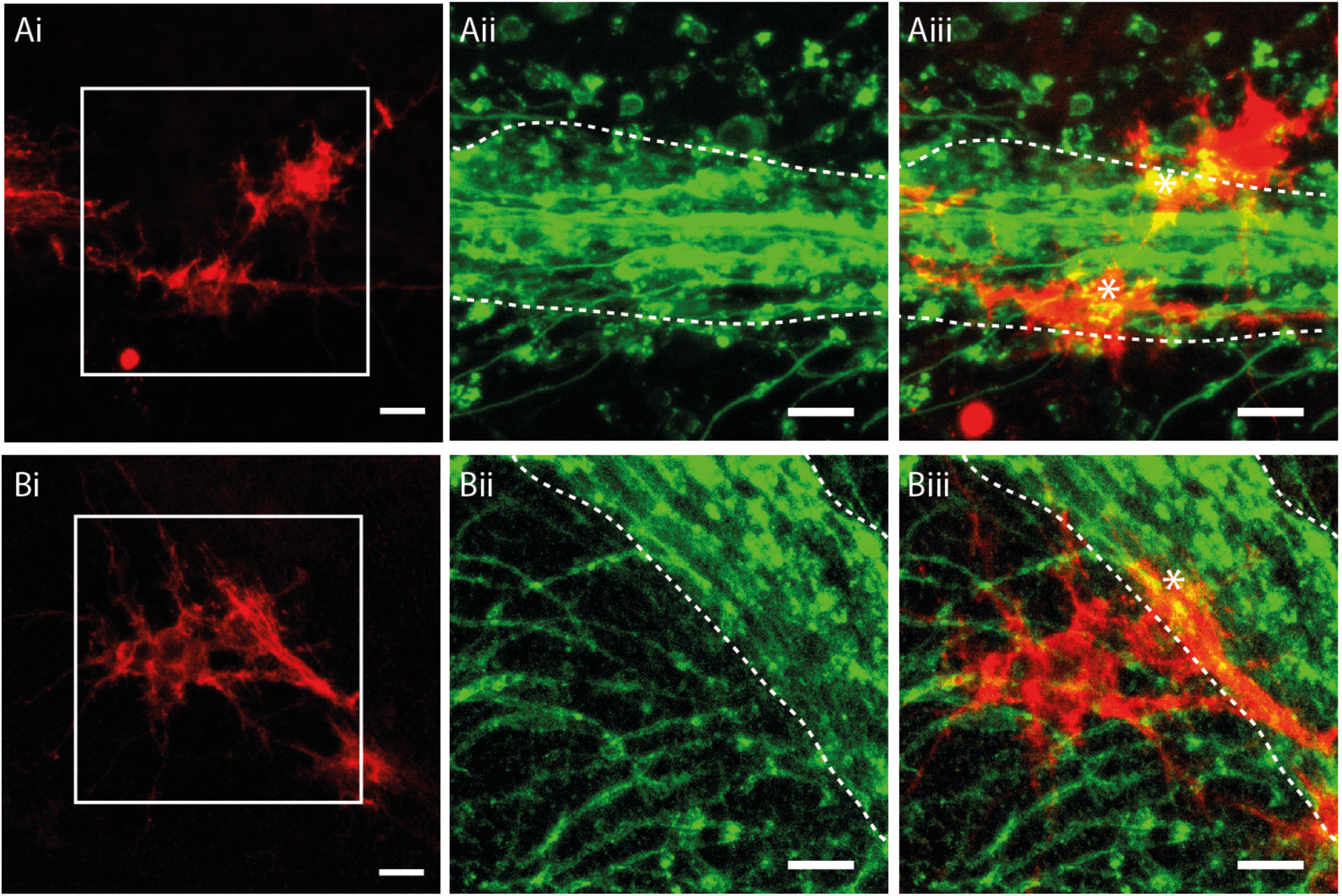
SFV transduced astrocytes are morphologically and antigenically distinct from OL. Ai, Bi. mCherry-f labelled glia in cerebellar white matter exhibiting complex finely branching process fields typical of astroglia. Aii, Bii. Enlarged view of the boxed areas shown in Ai/Bi showing the major white matter paths (dashed white lines) with MBP^+^ OL processes (green) branching into the granule cell layer. Aiii, Biii. Merged image showing mCherry-f fluorescence and MBP immunoreactivity from the same enlarged views shown in Aii/Bii. White dashed lines indicate major white matter paths. In contrast to OL (Fig. 1) SFV labelled astrocytes lack cellular processes aligned in parallel with the major white matter tracks, or linear segments of MBP/process localization. Astrocytes exhibit patches of MBP co-localization (*) suggesting contact with MBP^+^ profiles within cerebellar white matter. Scale bars in all panels 20 µm.

### Morphological and immunohistochemical features that distinguish labelled myelinating OL from astrocytes

The injection of SFV vectors into hippocampal slice cultures labells glia displaying a morphology typical of astrocytes [18, 24] with highly-ramified GFAP^+^ processes that are immunonegative for the OL marker RIP [18]. In addition to the OL described in Fig. 1, we also observed astrocyte-like glia following injection of SFV into cerebellar white matter (Fig. 2Ai, Bi). These cells were easily distinguished from OL by their distinct morphology, chiefly characterised by numerous finely branching processes that were not oriented in parallel with the major white matter paths in the folia (Fig. 2Ai, 2Bi). In addition, the processes of these cells lacked the linear segments of MBP localisation typical of OL (Fig. 2Aiii, 2Biii). In some cases small patches of MBP localization were present (Fig. 2Aiii, 2Biii), likely reflecting convergence between astrocytes and the processes of MBP^+^ OL, as has been reported previously for NG2^+^ glia [25]. Overall SFV-labelled astrocytes were easily distinguished from myelinating OL by their distinct appearance and the absence of linear stretches of MBP reactivity that characterized OL processes (Fig. 2Aiii, 2Biii).

### Analysis of _T_OL morphology and internodes

We quantified the number, length and branching of OL process arbors (see Supplementary Materials and Methods online) measured from 21 cells labelled by SFV-eGFP-f. Myelinating OL arbor morphology varied from a few longitudinally arranged process arbors (Fig 3A) to a complex collection of parallel-aligned longitudinally arranged processes (Fig 3B). On average OL in cerebellar white matter had 7.38 (± 0.64) process arbors (Fig. 3C) with an average length of 201 µm (± 23.63) and a maximum branch order of 7 (± 0.43) (Fig. 3D, 3E). In addition, _T_OL were found to support an average of 27.9 (± 2.5) branches across the network of their process arbors (data not shown). Using the criteria for internode identification (Methods) all of the OL examined contained process arbors with a mix of _NM_PB and _M_PB (Fig. 3Aiii, 3Biii) placing these cells at the _T_OL stage of maturation [20]. Quantification of _NM_PB and _M_PB revealed that the majority of process branches were non-myelinating (ratio _NM_PB/TB 0.83; ratio _M_PB/TB 0.17) (Fig. 3F, 3G). SFV labelled _T_OL had an average of 4.14 (± 0.43) internodes / cell (Fig. 3H), with a mean internode length of 103.8 µm (± 8.54) (Fig. 3I). _T_OL internodes were contained within an area extending an average of 36.5 µm (range 23.9 to 42.7 µm) perpendicular to the white matter path, and 17.33 µm (range 15.6 to 22 µm) in the Z axis forming a flattened tube-like space. The spatial arrangement of internodes was examined by analyzing the branch ordering of internode initiation sites, and quantifying the maximum number of internodes per _T_OL process arbor (Table 1, control study). Internodes initiated from branches located across a wide range of orders (range 1^st^ to 9^th^ order), while the number of internodes associated with each _T_OL process varied from a single internode (Fig.3 Aii, Aiii), to process arbors supporting up to 5 internodes interconnected by fine connecting branches (Fig. 3Biii, inset).

**Figure 3.**
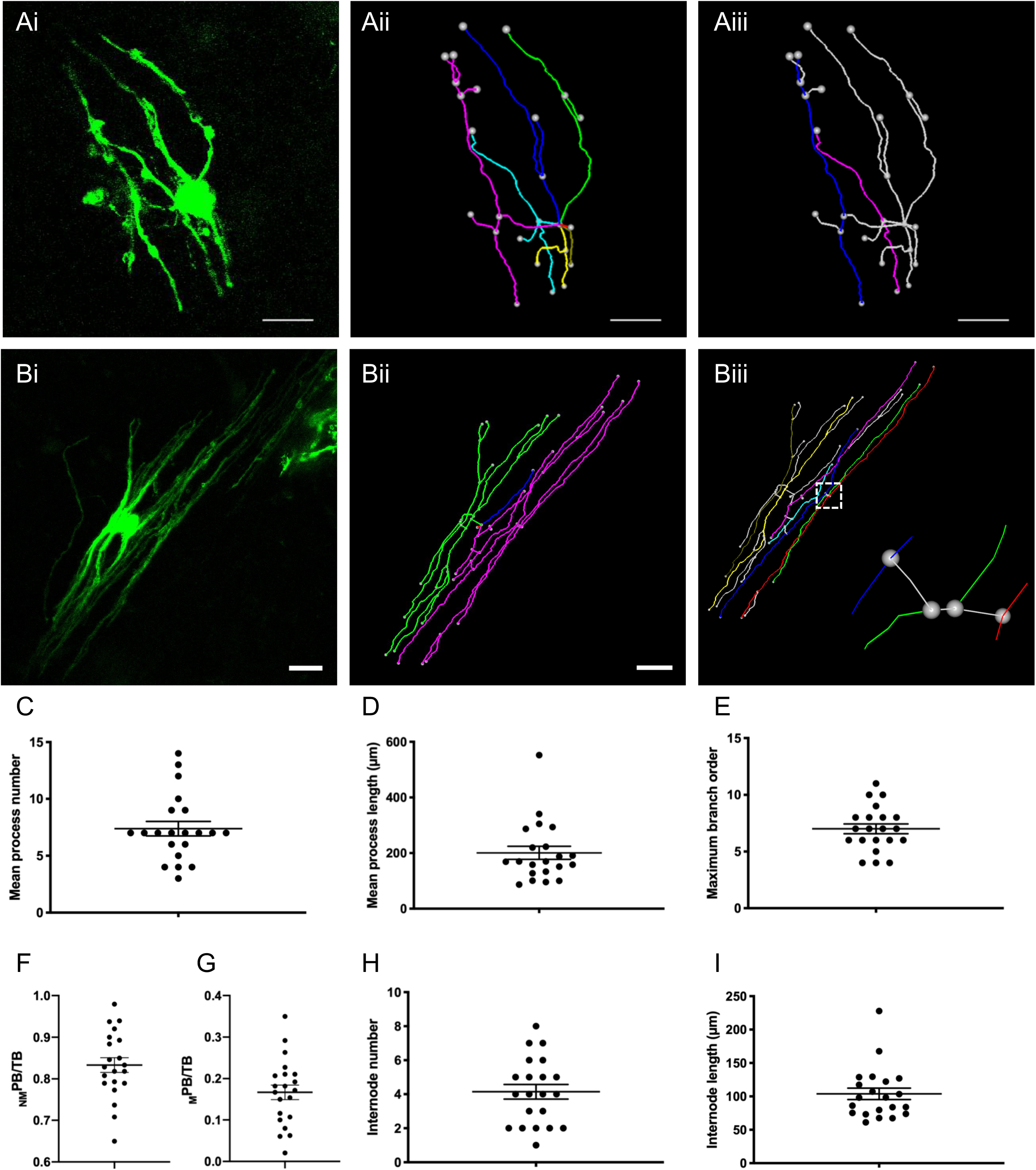
3-D reconstructions of SFV-labelled OL and quantification of OL process morphology. Ai. Maximum projection showing eGFP-f expression in a relatively simple white matter OL transduced with SFVA7(74). Aii. 3-D reconstruction of the cell shown in Ai. Colours (cyan, blue, green, yellow, purple, white) indicate six individual processes. Aiii. Reconstruction of the cell shown in Ai indicating position of two identified internodes (Blue, Purple). Note in this simple OL each internode initiates from a distinct processes. Bi. Maximum projection showing eGFP-f expression in a relatively complex OL. Bii. 3-D reconstruction for the cell shown in Bi. Colours (blue, green, purple) indicate three individual process arbors. Biii. Reconstruction of the cell shown in Bi indicating seven identified internodes (cyan, blue, green, yellow, purple, red, white). Note, individual processes (green and purple in Bii) support multiple internodes. Inset: Expanded view of area indicated by dashed box reveals the initiation of internodes from neighboring internodes via fine connecting processes (white segments). Scale bars in A and B 20 µm. C-G. Scatter plots displaying features of OL process morphology: mean process number per OL (7.38 ± 0.64) (C), mean process length (201 ± 26.63 µm) (D), maximum branch order reached per process (7 ± 0.43) (E), ratio _NM_PB/TB (0.83 ± 0.02) (F), and ratio _M_PB/TB (0.17 ± 0.02) (G). H-I. Mean internode number per cell 4.14 ± 0.43 (H) and internode length 103.8 ± 8.54 (I) derived from 21 SFV labelled OL.

**Table 1.** Morphology of _T_OL

|  | (1) Control study | (2) TTX study |  |  |
| --- | --- | --- | --- | --- |
|  |  | Control | TTX | *P* |
| **Process arbors** |  |  |  |  |
| Length ($\mu m$) | 201 ($\pm 23.63$) | 197 ($\pm 16.56$) | 169.82 ($\pm 11.46$) | 0.27 |
| Maximum branch order | 7 ($\pm 0.43$) | 7.2 ($\pm 0.50$) | 6.9 ($\pm 0.53$) | 0.68 |
| **Internodes** |  |  |  |  |
| Minimum order | 1.2 ($\pm 0.19$) | 1.1 ($\pm 8.51$) | 1.3 ($\pm 0.19$) | 0.85 |
| Maximum order | 4.76 ($\pm 0.50$) | 4.5 ($\pm 0.52$) | 3.5 ($\pm 0.38$) | 0.16 |
| Mean order | 2.64 ($\pm 0.23$) | 2.6 ($\pm 0.24$) | 2.2 ($\pm 0.22$) | 0.18 |
| $\bar{x}$ internodes / process | 2.2 ($\pm 0.26$) | 2.4 ($\pm 0.22$) | 1.9 ($\pm 0.17$) | 0.24 |
| Minimum and Maximum order refers to order of initiating branch. (1) relates to data shown in Figure 3, (2) relates to data shown in Figure 6. Values are Mean ± SEM. *P* values report comparison of Control and TTX data by Mann-Whitney *U* test. | | | | |

### Morphological analysis of OL after complement-mediated myelin injury

We examined the ability of the tracing method to detect changes in OL morphology in cerebellar slices exposed to a complement-mediated myelin injury [26] (see Supplementary Materials and Methods online). Consistent with an injury to OL and myelin, MBP staining of complement treated slices revealed discontinuous segments of myelin with a globular appearance, while control slices treated with complement alone, or an IgG isotype control, displayed intact MBP^+^ segments with a smooth appearance (see Supplementary Fig. S2 online). To determine how complement injury affected the morphology of OL SFV, vectors encoding an mCherryMBP fusion protein were injected into cerebellar white matter 24 h after the onset of the injury treatment. OL tropism for the mCherryMBP transgene was confirmed in slice cultures prepared from CNPase-GFP OL reporter mice [23], where it was targeted to _T_OL expressing the CNPase-GFP^+^ transgene (Fig. 4Ai-iii). Although the mCherryMBP label provided a strong signal that clearly illuminated the process arbors of transduced OL, the fluorescence exhibited a distribution that was distinct to the other SFV-delivered proteins, e.g. eGFPf (Fig. 1Bi, 1Ci), in being enriched at specific regions within the process arbors of _T_OL (Fig. 4Ai). Despite this punctate labelling, the mCherryMBP signal was suitable for tracing, with reconstructions from the non-injured control condition revealing numerous parallel aligned process arbors and putative internodes as expected of myelinating _T_OL (Fig. 4Aiv; see supplementary movie S2 online). In contrast, analysis of _T_OL in the complement injured slices showed clear signs of cellular damage characterized by the fragmentation of process arbors (Fig. 4Bi). This injury was evident in 3-D reconstructions, where intact process arbors were scarce, disconnected SFV labelled segments were frequent and internodes were not detected (Fig. 4Biv; see supplementary movie S3 online).

### Inhibition of neuronal activity reduces myelination in cerebellar white matter

To determine whether the SFV tracing method was sensitive to changes in myelination the morphology of OL was assessed after pharmacological blockade of neuronal activity. We have previously reported that sustained incubation in TTX for 48 h reduced myelination in cerebellar slice cultures [5]. Considering the variability in internode number and length obtained from SFV labelled OL (Fig. 3C, 3D), we examined the effect of a longer reduction in neuronal activity after incubation in TTX for 7 d. We first confirmed the effects of this treatment on axonal density and myelination by immunofluorescent staining for NF200 and MBP, respectively (Fig 5A, 5B) (see Supplementary Materials and Methods online). The averaged NF200 pixel area for slices treated with TTX was reduced compared to controls (Fig. 5C) (*P*<0.05) suggesting a reduction in the density of axons in TTX treated slices. MBP/NF200 co-localisation was also reduced in TTX-treated slices indicating a reduction of myelination (Fig. 4D) (*P*<0.001). Importantly, the reduction in myelination was independent of the effects on axonal density since co-localisation was normalised against axonal density (Methods).

**Figure 4.**
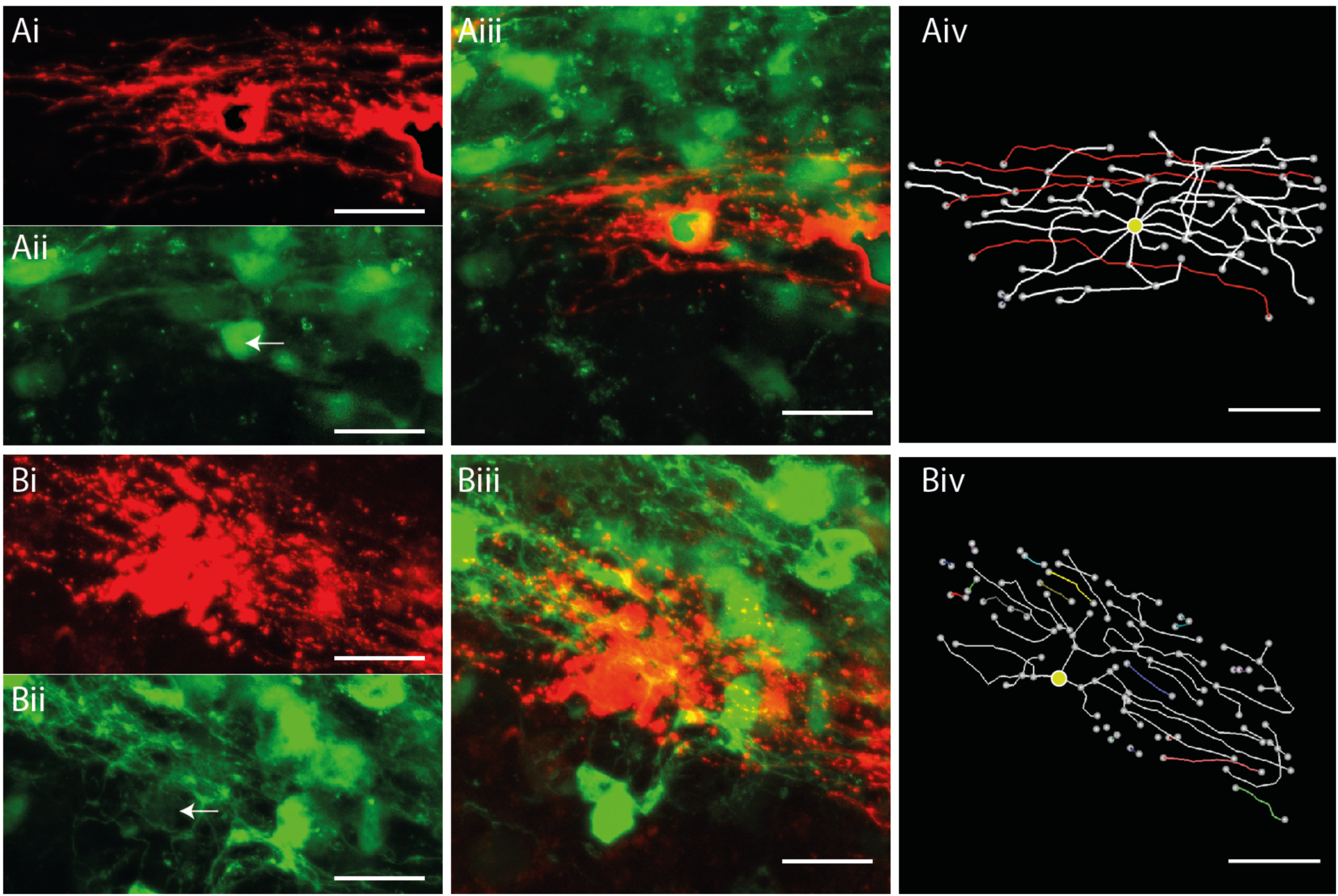
Complement mediated injury destroys internodes on SFV labelled OL. A. Reconstruction of an SFV labelled OL expressing the CNPase-GFP transgene in a complement control treated slice. Ai. mCherryMBP fluorescence reveals an OL with a typical myelinating morphology. Aii. CNPase-GFP signals imaged from the field shown in Ai (white arrow indicates position of the mCherry^+^ OL). Aiii. Merged image of the field shown in Ai-ii confirms co-localisation of mCherry (red) and CNPase-GFP (green) signals. Aiv. 3-D reconstruction and analysis reveals 4 internodes (red) on the OL shown in Ai-iii. B. Morphological analysis of an A774 labelled OL following complement injury. Bi. mCherry-f labelled OL with degenerating processes. Bii. CNPase-GFP signals imaged from the field shown in Bi. White arrow indicates position of mCherry^+^ OL shown in Bi. Biii. Merged image reveals localization of mCherry (red) and CNPase-GFP (green) signals. Biv. Reconstruction and analysis of the cell shown in Bi-iii reveals fragmented processes (coloured segments) and the absence of internodes. Scale bars in A and B 20 µm.

**Figure 5.**
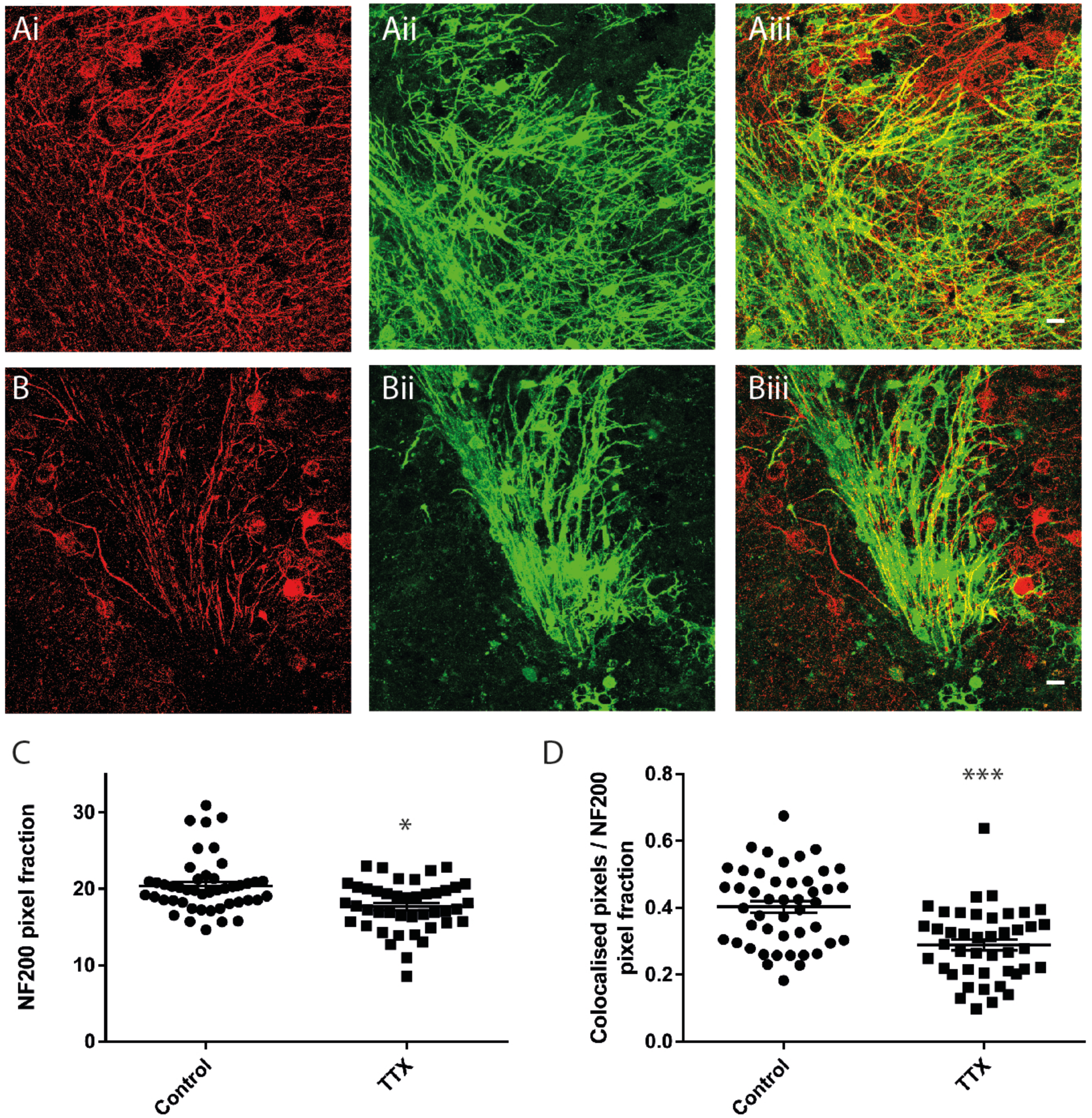
Sustained inhibition of neuronal activity reduces myelination of cerebellar white matter. A, B. Maximum projection images from representative control (A) and TTX (B) treated cerebellar slices showing immunofluorescent signals for anti-NF200 (Ai, Bi) anti-MBP (Aii, Bii), and co-localisations used to quantify myelination (Aiii, Biiii). C. Quantification of anti-NF200 signals. Mean NF200 pixel fraction is reduced by TTX (Control 20.4 ± 0.5, TTX 17.7 ± 0.5). D. Average MBP/NF200 ratios normalized against the NF200 signal are significantly reduced by TTX (Control 0.4 ± 0.02, TTX 0.29 ± 0.02). Scale bars in A and B 20 µm. * and *** Significance *P*<0.05 and *P*<0.001, respectively. Data expressed as means ± SEM.

### Neuronal activity influences _T_OL process arbor morphology

Having confirmed that a 7-d incubation with TTX reduced myelination in cerebellar slice cultures, we examined whether sustained inhibition of neuronal activity altered the morphology of SFV labelled _T_OL. Representative 2-D reconstructions of _T_OL from control and TTX treated slices are displayed in Figure 6A and 6B, and 3-D movies of these cells can be viewed in supplementary movies S4 and S5 (available online), respectively. TTX treatment increased the mean number of process arbors per _T_OL (*P*<0.05) (Fig. 6C), and the branching of these process arbors (described by value TB) (*P*<0.05) (Fig. 6D). In contrast, TTX treatment did not alter the average length of process arbors (Table 1, TTX study), or the maximum branch order attained by these processes (Table 1, TTX study). In agreement with the data on myelination (Fig. 5D), TTX treatment decreased the length of internodes (*P*<0.05) (Fig. 6E). Similarly, internode number tended to be lower in TTX treated _T_OL, although this decrease did not reach statistical significance (*P*=0.08) (Fig. 6F). To explore the morphology of _T_OL further we analysed the ratios _NM_PB/TB and _M_PB/TB on control and TTX treated _T_OL. TTX treatment increased the ratio _NM_PB/TB (Fig. 6G) (*P*<0.001), while decreasing the ratio _M_PB/TB (Fig. 6H) (*P*<0.0001) implying that blockade of neuronal activity decreases the proportion of myelinating branches. We also examined whether neuronal activity influenced the spatial arrangement of internodes by quantifying the rank order of internode initiation sites. TTX had no effect on the spatial arrangement of internode initiation sites, with neither the minimum, maximum or average rank of initiation being altered between control and TTX treated _T_OL (Table 1, TTX study). Similarly, TTX treatment had no effect on the maximum number of internodes per process arbor (Table 1, TTX study).

**Figure 6.**
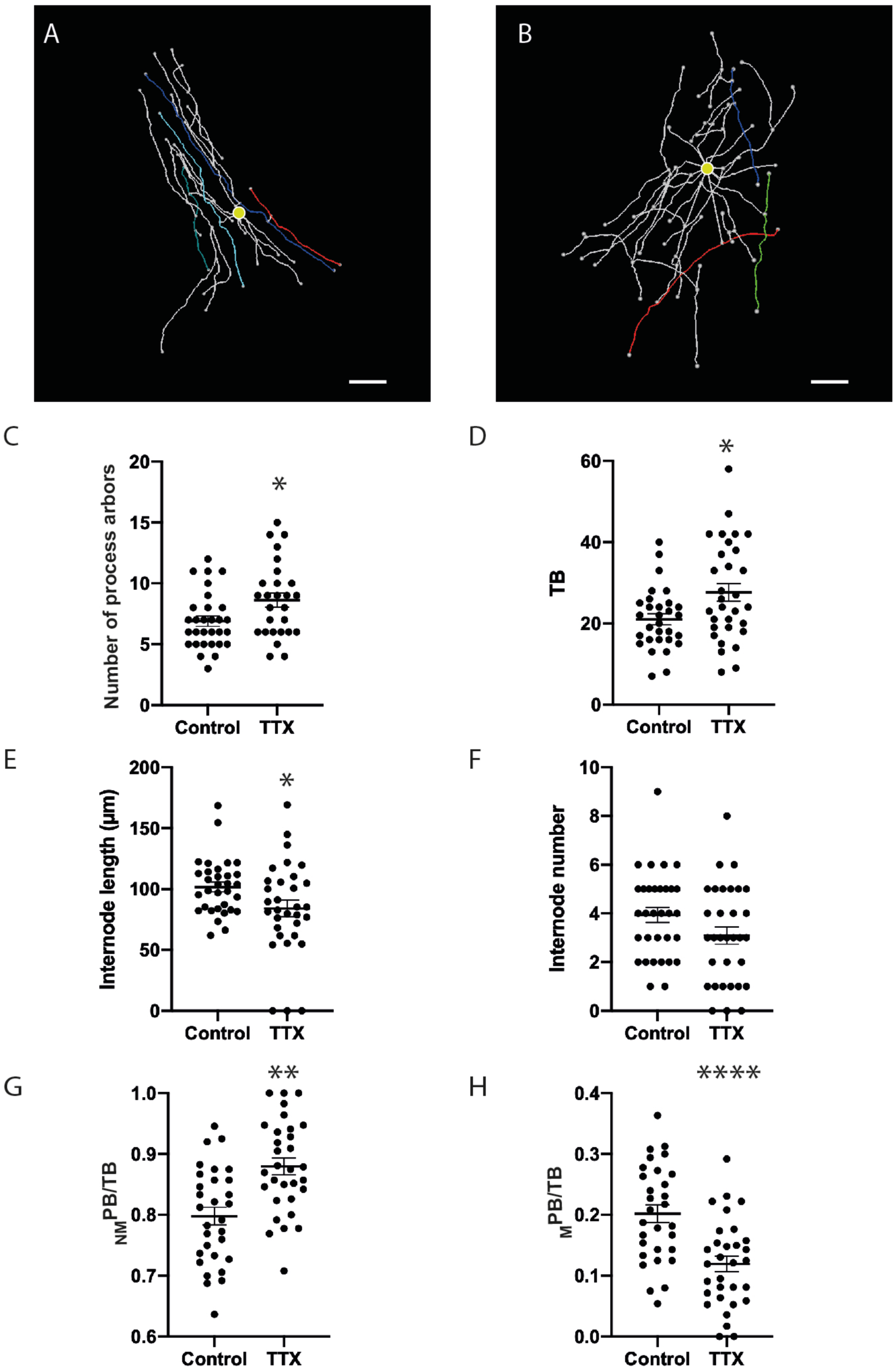
Sustained inhibition of neuronal activity alters OL morphology. A-B. Representative 3-D reconstructions of OL from control (A) and TTX (B) treated cerebellar slices showing internodes selected for quantification (A: cyan, blue, green, red; B: blue, green, red) and cell bodies (yellow circles). C-D. TTX treatment increased the number of processes arbors per OL (C) (Control 6.9 ± 0.4, TTX 8.63 ± 0.6) and the total number of branches (D) (Control 21.03 ± 1.38, TTX 27.65 ± 2.18). E-F. TTX treatment reduced the mean internode length of OL (E) (Control 101.8 µm ± 4.1, TTX 85.4 µm ± 6.7), but not the absolute number of internodes (F) (Control 4.2 ± 0.3, TTX 3.2 ± 0.4). G-H. TTX treatment induced complementary changes in the ratio _NM_PB/TB (G) (Control 0.8 ± 0.01, TTX 0.88 ± 0.01) and _M_PB/TB (H) (Control 0.2 ± 0.01, TTX 0.11 ± 0.01). Scale bars in A&B 20 µm. * Significance *P*<0.05, ** Significance *P*<0.01, **** Significance *P*<0.0001. Data expressed as means ± SEM.

## DISCUSSION

We have used SFV labelling to develop new findings on the activity-dependent maturation of _T_OL, and their response to complement-mediated injury. We show that SFV labelling in neonatal cerebellar slice cultures provides a platform for analyzing the effects of demyelinating injuries and altered axonal activity on _T_OL process morphology, and internode number and length. This approach has several useful features. First, the use of organotypic slice cultures allows pharmacological modulation of neuronal function, and OL viability, within a system in which the *in vivo* arrangement of neuronal and glial cells is well preserved. Second, the slice cultures provide an opportunity to image OL in deep lying white matter structures, such as those of the cerebellum, that are beyond the reach of current *in vivo* imaging methods [17]. Third, SFV labelled slices are amenable to long-term live-imaging [19], thus they may be exploited in imaging-based screening studies focused on dissecting the mechanisms regulating the effects of local circuit activity and CNS injury on OL and internode morphology.

SFV labelling in the white matter of cerebellar slice cultures revealed cells whose morphology and antigenic profile identified them as differentiated OL. The use of membrane tethered FPs enabled accurate tracing and analysis of OL morphology revealing cells with complex process arbors incorporating multiple branch points, and giving rise to both _NM_PB and _M_PB, the latter of which were identified as thickened longitudinal segments of membrane attached to the cell *via* fine connecting branches. This combination of complex process fields with both _NM_PB and _M_PB identifies these cells as _T_OL, a developmental stage positioned between premyelinating and _M_OL [19, 20]. In support of this classification, SFV labelled-OL share morphological characteristics with ‘immature OL’ labelled in rat optic nerves [27, 28]. Moreover, the present work, and that of previous studies describing _T_OL/immature OL phenoptypes [20, 27, 28], did so in tissues of a similar age (e.g. postnatal CNS tissues P5-14), thus the OL labelled by SFV injections in cerebellar white matter are of an intermediate maturity between pre-myelinating and _M_OL.

Although several studies have described the morphology of _T_OL [19, 20], the present work is the first to describe the spatial distribution of internode initiations with respect to branch ordering. Initiation points were not restricted to the primary branch, but occurred in a wide range of orders.

These findings imply that the potential for internode initiation is present at all orders, and that transport mechanisms for myelin basic protein mRNA, which undergoes local translation at sites of myelination [29] (reviewed in [17]), are capable of delivering MBP mRNA to all orders of a process arbor. In agreement with previous work from neonatal medulla [20], cerebellar _T_OL process arbors supported up to 5 internodes interconnected by fine process extensions. Internodal initiation sites have not been reported from _M_OL in adult CNS tissues [15, 28, 30] suggesting that they may reflect a transient developmental phenomenon. Future work involving long-term live imaging can address this question directly by tracking the fate of these internodes.

SFV labelled _T_OL possessed an average of 5 internodes (range 1 to 8), lower than comparable data obtained *in vivo*, which consistently report around 20 internodes per OL [27, 28, 30, 31]. These *in vivo* studies have largely been pursued in the adult optic nerve where _M_OL are expected to predominate, thus the lower numbers of internodes reported here may reflect the immature nature of the _T_OL under study. Interestingly, measurements from a number of different adult CNS tissues revealed that OL in the cerebellum possess the fewest internodes [32], although this number is still 4-fold greater than that reported in the present work.

The internodal lengths measured from _T_OL in cerebellar slices (mean 106 µm) are in good agreement with those measured from cerebellar OL *in vivo* [32] where a distribution centered around ∼100 µm is observed. Chong et al. [32] also used sparse OL labelling with a membrane targeted GFP to accurately resolve the 3-D morphology of OL and their internodes. The similarity in these approaches, and the resulting data, provide confidence in the methods we have used to identify and quantify internodes in _T_OL. Our *ex vivo* measurements of cerebellar internode lengths are considerably longer than those obtained from neocortical slice cultures (mean 49µm,[16]). Importantly, cortical internodes measured *in vivo* fall in a similar range to those measured *ex vivo* (∼50 to 60 µm [32]) indicating that regional variations in internode length observed *in vivo* are preserved in slice cultures.

Our results provided new insights into complement-mediated injury in OL. 3-D analysis of SFV labelled _T_OL enabled us to resolve the degeneration and loss of process arbors in slice cultures subjected to complement-mediated demyelinating injury. A significant outcome for this work was the absence of identifiable internodes after complement-induced injury, an observation that corresponded with a degradation in the integrity of myelin segments visualized by MBP staining. Previous studies imaging complement injury in slice cultures have used slices from OL reporter mice where transgene expression is present in all OL [26, 33]. Thus parameters, such as process arbor morphology and internode number or length, could not be resolved at the single cell level. This work is therefore the first to provide a description of complement-induced OL injury at the level of single OL. These advances are important since auto-antibodies directed against myelin antigens, for example MOG (the myelin epitope targeted in the present work), are associated with OL injury and demyelination in Multiple-Sclerosis [34, 35]. Moreover, cerebellar slice cultures have recently been used to investigate the pathogenicity of recombinant auto-antibodies derived from the CSF of MS patients [33], and are gaining recognition as a useful tool for mechanistic studies into human demyelinating disease. In this context, live-imaging studies of SFV labelled OL could yield new information on the time-course OL injury, and be adapted to provide a platform for screening oligo-protective compounds.

The analysis of SFV labelled _T_OL enabled the influence of neuronal activity to be studied during the early stages of myelination. Sustained treatment with TTX diminished the localization of MBP signals on cerebellar axons agreeing with our earlier work where a briefer incubation produced similar effects on myelination [5]. Analysis of SFV labelling after TTX treatment allowed us to correlate changes in the morphology of _T_OL with reduced myelination. Blockade of neuronal activity increased the number of process arbors and the total number of branches they supported. In addition, TTX reduced the proportion of branches supporting internodes (MB/TB), a ratio that is expected to increase as OL differentiate into a fully mature myelinating phenotype in which all processes support internodes [20]. These observations from TTX treated _T_OL imply that neuronal activity guides the morphological differentiation of OL so that growth is channeled towards the production of internodes on arbors receiving activity-dependent cues from target axons. In contrast, when these cues are absent, growth is invested in the generation of additional process arbors and non-myelinating branches at the expense of myelinating segments. In agreement with this idea, TTX treatment increased the ratio _NM_PB/TB, indicating that _T_OL make a smaller investment in the production of internodes. These insights on the morphological response of myelinating OL complement earlier results revealing reduced myelination after treatments with TTX [4–6]. They also partially match *in vivo* findings from a social isolation paradigm where reductions in myelination are detected in the medial prefrontal cortex (mPFC) [8]. Here, a post-weaning social deprivation paradigm designed to reduce neuronal activity, decreased the number of internodes produced by mPFC OL. However, contrary to the present results, social isolation induced a decrease in the number of OL processes arbors, and the complexity of their branching. Differences in experimental approaches may explain these contrasting effects. TTX, as used here, will entirely block the activity of excitatory and inhibitory (GABAergic) neurons, while social isolation is likely to produce more nuanced effects that could differentially effect excitatory and inhibitory circuits. In this regard, recent work from neocortical slice cultures has uncovered an involvement of endogenous GABAergic signaling in cortical myelination [16], in which blockade of GABA-A receptors increased the number of OL and boosted myelination. Interestingly, TTX prevented these changes, but had no effect on OL numbers when applied alone suggesting that GABA influences neocortical OL through a combination of activity-dependent and independent mechanisms. These findings contrast clearly with data from the cerebellum in which TTX promoted both OPC proliferation and differentiation [5]. Thus, clear differences exist in the mechanisms regulating OL populations in the cerebellum and neocortex which may explain the differing morphological responses shown by OL in these regions.

TTX treatment reduced the length of internodes produced by cerebellar _T_OL. This finding agrees with *in vivo* data from mouse optic nerve where monocular deprivation, or the genetic inhibition of vesicle release, lead to reductions in internodal length [36]. These findings from cerebellar slices, and optic nerve, suggest the involvement of an activity-dependent axonal signal during the elongation of myelin sheaths. However, as suggested by Hamilton et al. [16], reductions in internodal length may also reflect the outcome of increased competition among OL for a limited number of target axons. Indeed, blockade of neuronal activity in both systems produces an increase in the differentiation of OL [5], and in cerebellar slices this may be accompanied by a small reduction in axonal density (Fig. 5). The screening of activity-dependent signaling molecules in *ex vivo* slice cultures would help to determine if neuronal activity exerts distinct actions on OL differentiation and internode length.

Global disruption of axonal neurotransmission via expression of a tetnus neurotoxin (TeNT) reduces both myelination and the number of internodes formed per OL in larval zebrafish [37]. However, silencing of global axonal activity by TeNT expression has no effect on internode length [37]. Interestingly, another study in zebrafish detected a reduction in internode length when activity was selectively silenced in a subset of *phox2b*^+^ spinal cord axons, but not when activity was silenced globally with TTX[38]. These discrepancies may arise through a competition-based interaction between adjacent axons since global suppression of activity by TTX prevented the reduction in internode length in TeNT expressing in *phox2b*^+^ axons. Developmental differences may also account for the discrepancy between findings from Zebrafish and the present study since the former data were obtained from larval CNS where newly emerging internodes are of a length (∼10µm) that would have been excluded from the present study (Methods). Indeed, the average lengths of these myelin segments are significantly shorter (∼ 10µm) than those examined in the mammalian CNS (∼100µm). Whether such activity dependent competition-based interactions operate at later stages of OL development in fish, or in the mammalian CNS remains to be determined.

In conclusion, the morphological analysis of SFV-labelled slice cultures enabled us to detect structural changes in individual cerebellar _T_OL including the early stages of OL degeneration during complement-mediated myelin injury and more subtle morphological changes resulting from the blockade of neuronal activity that included the growth of additional process arbors, and a reduction in the abundance and length of myelin internodes. Live-imaging studies based on this approach could be used to refine our understanding of OL injury and explore the neurochemical mechanisms supporting myelin plasticity.

## METHODS

### Animals

Organotypic cerebellar slice cultures were obtained from 50% C57BL/6J 50% CBA/CaCrl mice bred at the University of Warwick, or from either C57BL/6J mice or CNPase-GFP mice [23] bred at the University of Birmingham. CNPase-GFP mice, originally produced in the laboratory of Dr Vitorrio Gallo (Children’s National Medical Center, Washington, D.C.), were rederived from frozen embryos cryo-stored by Professor Attila Sik (University of Birmingham). Mice were killed according to humane methods proscribed in Schedule 1 of the Animals (Scientific Procedures) Act 1986.

### Organotypic slice cultures

Slice cultures were prepared and cultured as described in [5]. In brief, parasagittal vibratome slices (350µm) cut from the cerebellum of P8-11 mice were cultured on small pieces of confetti membrane (Merk Millipore, MA, USA) placed on top of culture inserts (Merk Millipore). Slices were cultured in 1 mL of culture media containing 50% MEM with Glutamax (Life Technologies Ltd, Paisley, UK), 23% EBSS (Life Technologies Ltd, Paisley, UK), 25% horse serum (Sigma-Aldrich Company Ltd, Dorset, UK), 6.5 mg/mL glucose, and 1250 units penicillin / streptomycin. Slices were maintained in a humidified incubator (37°C, 5% CO2) for 7 to 10d with media changes every 2–3d. In some experiments, slices were incubated in TTX (1µM, Ascent Scientific, Cambridge, UK) to block neuronal activity. TTX incubations began at 0 DIV and were continued for 7d with the medium/TTX replaced every 3d.

### 3-D Reconstruction of SFV-labelled OL

3-D reconstructions were prepared using NeuronStudio software (Computational Neurobiology and Imaging Center, Icahn School of Medicine at Mount Sanai;[39]). Cells were traced manually by viewing images in three-dimensions using the Stack Viewer mode and the resulting reconstructions exported as swc files. For an illustration of the tracing method and the resulting 3-D reconstruction, see supplementary movie S1. For methods used to analyse the morphology of OL and internodes see Supplementary Materials and Methods (online).

### Statistical analysis

Normal distributions in each data set were evaluated by the Shapiro-Wilk test. Two-group comparisons of normally distributed data were performed with *t* tests, while non-normal data were analysed using the Mann-Whitney *U* test. Statistical calculations were performed using Prism 7.0 (Graphpad Software, El Camino Real, CA, USA). Reliable differences between groups were accepted at a two-tailed *P*α<0.05. All data are presented as mean ± SEM.

Additional methods can be viewed in the Supplementary Materials and Methods file.

## SUPPLEMENTARY INFORMATION

See accompanying file for Supplementary Materials and Methods and supplementary data.

## ACKNOWLEDGMENTS

This work was supported by a fellowship to D.F. from the Birmingham Science City Research Alliance funded by the Higher Education Funding Council for England (HEFCE), the People Programme (Marie Curie Actions) of the European Union’s Seventh Framework Programme (CIG 294051), and the University of Birmingham’s Wellcome Trust Institutional Strategic Support Fund (2014 round). We thank Dr Zubair Ahmed for his useful comments on the manuscript, and Professor Pranab Das (University of Birmingham) for discussions and advice on complement-mediated injury.

## AUTHOR CONTRIBUTIONS

Concept and design of the study: D.F.; design and generation of SFVA7(74)mCherry-MBP vector: S.M.R. and D.F.; data acquisition D.F., data analysis: E.T. and D.F.; drafting of manuscript and figures: M.B. and D.F. All authors read and approved the submitted version of the manuscript.

