## Supplementary Method and Supplementary Data for "A morphological analysis of activity-dependent myelination and myelin injury in transitional oligodendrocytes"

### Supplementary Materials and Methods

#### Semliki Forest Virus preparation

Recombinant Semliki Forest Virus vectors of subtype A7(74) (SFVA7(74) were used to transduce OL with the following transgenes: Cytosolic mCherry, farnesylated mCherry (mCherry-f), farnesylated eGFP (GFP-f), and an mCherry-myelin basic protein fusion (mCherry-MBP). cDNA encoding SFVA7(74)-mCherry and GFP-f were kindly provided by Markus Ehrengruber (Kantonsschule Hohe Promenade, Zurich). The SFVA7(74)-mCherry-f vector was constructed by inserting an mCherry-f sequence (obtained from Dr Yann Bernardinelli [22] into the multiple cloning site of an empty SFVA7(74) cDNA (obtained from Dr Markus Ehrengruber). The sequence for the mCherry-MBP fusion was provided by Professor Mikael Simons (Max Planck Institute of Experimental Medicine, Gottingen, Germany;[23]). An SFVA7(74) cDNA encoding this mCherry-MBP sequence was synthesized commercially (Genewiz LLC, South Plainfield, NJ, USA). Infectious particles were generated from these SFVA7(74) vectors in BHK cells using the methods described by Ehrengruber, Schlesinger [24]. Un-purified viral preparation were then aliquoted and stored at -80 °C.

#### Viral infection of slice cultures

To infect white matter OL SFV particles were injected directly into the white matter. Aliquots of the virus where thawed and, except when stated, diluted ten-fold in sterile injection solution. Injection solution consisted of DMEM with elevated $MgCl_2$ (10 mM), TTX (0.5 μM) and penicillin/streptomycin (125 units/mL). The final concentration of SFV particles ranged from from $10^5$ to $10^7$ infectious units /mL. Injections were made using glass micropipettes fabricated from standard glass capillaries (World Precision Instruments, Sarasota, FL, USA). Pipettes were backfilled with 20μL of diluted viral stock and mounted into an air-tight electrode holder attached to a micromanipulator. The electrode holder was then connected to a 1mL syringe via plastic tubing with a 3-way valve located between the syringe and the tubing to permit charging of the syringe without disturbance of the pipette contents. Slice cultures attached to confetti were transferred to the stage of an upright microscope and submerged with 1 mL of pre-warmed (37°C) injection solution. Slices were observed with a 10x or 20x objective and a monochrome CCD camera (Watec Incorporated, Newburgh, NY, USA) attached to the microscope. Injection pipettes were positioned slightly above the surface of the white matter and a small amount of pressure was applied to ensure positive ejection of viral solution. Having been lowered approximately 10-20 µm into the tissue with zero pressure, a small amount of positive pressure (2 seconds) was applied to eject viral solution. Positive ejection of virus was readily confirmed by a slight swelling of the targeted tissue. Typically, multiple injections were made along the long axis of each folia within the slice, with an average time to inject a slice of approximately 5 minutes. Following injection, slices were returned to a sterile culture hood where they were rinsed 1x in 2 mL of pre-warmed (37°C) wash media (DMEM containing penicillin/streptomycin (125 units/mL)). After rinsing slice cultures were returned to the culture well and incubated for a further 15 to 24 hours to allow expression of transgenes. In some experiments SFVA7(74)mCherry-f was applied directly to the surface of the slice using the method described by Fannon, Tarmier [5].

**Complement injury model**

CNPase-GFP slice cultures were injured using a protocol adapted from methods described by Harrer et al. [25]. Briefly, slices were incubated in culture medium supplemented with 10% baby rabbit complement (VH Bio Ltd., Gateshead, UK) and monoclonal anti-MOG (20 µg/mL, MAB 5680, Millipore (U.K.) Ltd., Watford, UK) at 37°C. Control slices were incubated in culture medium containing 10% baby rabbit complement. 24 hours after the start of the treatments slices were injected with SFVA7(74)mCherry-MBP as described above, and then returned to the treatment wells for a further 24 hours. Overall slices were incubated in complement +/- anti-MOG for 48 h, after which they were fixed in 4% paraformaldehyde (PFA) and mounted on slides. The same protocol was used to injury wild type slice cultures in experiments using MBP immunohistochemistry. In additional control experiments slice cultures were incubated in culture medium containing 10% baby rabbit complement plus mouse isotype control IgG (20 µg/mL, Sigma Aldrich, Cat# M5284).

**Immunohistochemistry and analysis of myelination**

Immunoflourescent staining of slice cultures was performed using the methods described by Fannon, Tarmier [5]. Briefly, slices were fixed in 4% PFA overnight at 4°C, washed four times in phosphate buffered solution (PBS), blocked and permeabilised at room temperature in PBS containing 10% normal goat serum and 0.2% triton X-100 (blocking solution), then incubated overnight at 4°C in primary antibodies diluted in blocking solution. The next day slices were rinsed in PBS four times, then incubated in secondary antibodies diluted in blocking solution for 4-5 hours at room temperature. Finally, slices were washed in PBS and mounted beneath glass coverslips with aqueous mounting media (Immu-Mount, Thermo Fisher Scientific, Paisley, UK). Myelin proteins were detected with rat-anti MBP (1:200, Merk Millipore, MAB386) and mouse anti-MOG (1:200, described above). Axonal neurofilament was localised with chicken anti-NF200 (1:10,000, Abcam, ab72996), and astrocytes with mouse anti-GFAP (1:200, Sigma Aldrich, G3893). Primary antibodies were visualised with Alexa Fluor® (AF) conjugated Goat IgG (Invitrogen, San Diego, CA, USA) developed against the appropriate species. Myelination was quantified by examining colocalisation of myelin (anti-MBP) and axonal (anti-NF200) immunoreactivity using the methods described by Fannon, Tarmier [5].

**Confocal imaging**

In some experiments (Fig. 1A-B, Fig. 3, Fig. 5) SFV confocal imaging was performed using a laser-based Leica TCS SP5 confocal microscope (Leica Microsystems, Wetzlar, Germany). Using this system image stacks (1024 by 1024 pixels, 8-16 line average, 0.2 μm steps) were acquired with a 40x 0.8 N.A. water immersion objective. In other experiments (Fig. 1C-D, Fig. 2, Fig. 4, Fig. 6) image stacks were acquired using a non-laser based Zeiss Axiovert microscope (Zeiss, Oberkochen, Germany) equipped with a metal halide light source and a differential spinning disk module (Revolution DSD, Andor Technology plc, Belfast, UK). On this setup image stacks (630 by 1005 pixels, 1 μm steps) were acquired with either a 20x 0.5 N.A. air objective, or a 40x 1.3 N.A. oil immersion objective.

**Analysis of OL reconstructions**

Reconstructions, consisting of sets of nodes and interconnecting segments (see Supplementary Fig. S3), were viewed and analysed in Amira 5.6 (FEI Company, Hillboro, Oregon, USA) using the Spatial Graph module. OL morphology was investigated by quantifying the number and length of process arbors, with processes defined as the entire collection of segments and nodes arising from a common segment connected to the cell body (Supplementary Fig. S3A, S3B). Process arbor branching was determined as described below (see myelinating status of process branches). Branch rank orders were determined for each segment using an automated function that assigns rank orders based on the spatial relationship to the cell body (root function in Amira), such that first order segments were directly connected to the root, and second order processes arise from the first order segments *etc* (Supplementary Fig. S3C). Processes were then identified using Amira's sub-graph function (Supplementary Fig. S3B), and the maximum branch order and average length (total length of all branches on each process arbor) calculated for each process arbor. These data were then averaged for all processes detected on the cell to produce single data points per cell. Putative internodes were identified by inspection of both the confocal stack and 3-D reconstruction using the following criteria: thickened $FP^+$ processes initiating at T-junctions forming from fine connecting branches. The majority of internodes in the CNS are reported to exhibit lengths ≥ 50 µm[27, 28]. Therefore, to exclude non-myelinating processes only thickened segments (as viewed in the confocal stack) initiating from a T junction and exhibiting a length ≥ 50 µm were annotated as internodes. Internodes were annotated within the Amira reconstructions by creating label groups containing individual nodes and segments associated with each internode. This operation allowed the number and length of each internode to be quantified by Amira. The spatial arrangement of internodes was examined by analyzing the branch order of the fine connecting branch giving rise to each internode, and by quantifying the average of the maximum number of internodes per OL process. The myelinating status of process branches (${}_{NM}PB$ and ${}_{M}PB$) was quantified as follows (Supplementary Fig. S4): First, ${}_{M}PB$ values were derived directly from the count of internodes (red segments in Supplementary Fig. S4) for each cell (number of internodes = ${}_{M}PB$). Second, values for ${}_{NM}PB$ were computed using the formula ${}_{NM}PB = TN - ({}_{M}PB*2)$, where TN (Terminal Nodes) represents all nodes with no further connecting segments. The function ${}_{M}PB*2$ was used since each internode contains two

TN within the reconstruction. Finally, the total of all branches (TB, $_{NM}PB + _{M}PB$) was used to describe process arbor branching, and as a denominator for the ratios $_{NM}PB/TB$ and $_{M}PB/TB$.

**SUPPLEMENTARY FIGURES**

**Supplementary Figure S1. SFV vectors transduce CNPase-GFP expressing oligodendrocytes.** SFV vector transduces CNPase-GFP expressing myelinating oligodendrocytes. A. Maximum projection image showing an SFV labelled white matter cells with a typical myelinating morphology. B. GFP fluorescence from the field shown in A. C. Merge image indicates expression of the CNPase-GFP transgene within coma of SFV labelled cell. Scale bars 20 μm.

**Supplementary Figure S2.** Complement-mediated myelin injury in cerebellar slice cultures. A&B. Representative immunofluorescent staining for MBP in complement treated control slices reveals a high density of continuous $MBP^{+}$ segments. C. MBP staining from a control slice treated with complement plus an istotype control IgG. D. Characteristic injury to $MBP^{+}$ segments in a slice treated with complement plus anti-MOG. Note the abundance of of globular discontinuous MBP+ figures (arrows). Scale bars in all panels 20 μm.

**Supplementary Figure S3.** Identification of process arbors and branch orders from cell reconstructions. A. Maximum projection showing eGFP-f expression in a white matter OL transduced with SFV. B. 3-D reconstruction of the cell shown in A consisting of a set of nodes (coloured dots) with interconnecting branches. Note reconstruction has 8 individual process arobrs identified by distinct colours. C. Alternative display of the reconstruction shown in B with individual branches coloured according to their rank ordering: 1$^{st}$ order - red; 2$^{nd}$ order - cyan; 3$^{rd}$ order - green; 4$^{th}$ order - blue; 5$^{th}$ order - grey; 6$^{th}$ order - teal. Inset: Expanded view of area indicated by dashed box highlights rank ordering of branches on a single process arbor (colour scheme as above). Scale bars in A and B 20 μm.

**Supplementary Figure S4.** Procedures used to quantify $_{M}$PB and $_{NM}$PB. A. Maximum projection image showing an SFV labelled $_{T}$OL in cerebellar white matter. B. 3-D reconstruction of the cell shown in A with colour coded nodes (terminal nodes are colour coded for illustrative purposes only). Blue terminal nodes are associated with $_{NM}$PB; Yellow nodes are associated with $_{M}$PB. C. Rotation and zoom of the reconstruction shown in B provides a clear view of myelinating and non-myelinated branches. Total value of $_{M}$PB is derived from the number of internodes (red segments capped by yellow terminal nodes, 3 in this example). $_{NM}$PB were calculated by subtracting $_{M}$PB*2 from the total count of terminal nodes on the cell (21 in this example), thus $_{NM}$PB = 21-(3*2) = 15. Scale bars in A, B and C 20 μm.

**SUPPLEMENTARY MOVIES**

Supplementary movie files are available as separate files from Biorxiv.org.

**Supplementary movie S1.** Movie illustrating tracing of OL processes on an SFV labelled OL. Progress through consecutive confocal sections through the Z axis reveals tracing of OL processes. Eight individual processes are indicated in distinct colours. Upon completion of the Z scan the 3-D arrangement of processes in the reconstruction are displayed by a full rotation in the Y axis.

**Supplementary movie S2.** Movie showing 3-D structure and quantified internodes (yellow) for the non-injured complement control treated OL reconstruction depicted in Figure 4A. Rotations in the Y and X axis reveal the spatial arrangement of processes and internodes in the reconstruction.

**Supplementary movie S3.** Movie illustrating fragmentation of processes for the injured complement /anti-MOG treated OL reconstruction depicted in Figure 4B. The spatial arrangement of process fragments (colours) are revealed in sequential rotations of the reconstruction in the Y and X axis.

**Supplementary movie S4.** Movie showing 3-D structure and quantified internodes (colours) for the control OL reconstruction depicted and quantified in Figure 6A. Consecutive rotations in the Y and X axis are shown to provide a complete view of the 3-D arrangement of processes and internodes in the reconstruction.

**Supplementary movie S5.** Movie illustrating the 3-D structure and quantified internodes (colours) for the TTX treated OL reconstruction depicted and quantified in Figure 6B. A sequence of full rotations in the Y and X axis provide a complete view of the spatial relationship of processes and internodes in the reconstruction.

Figure S1

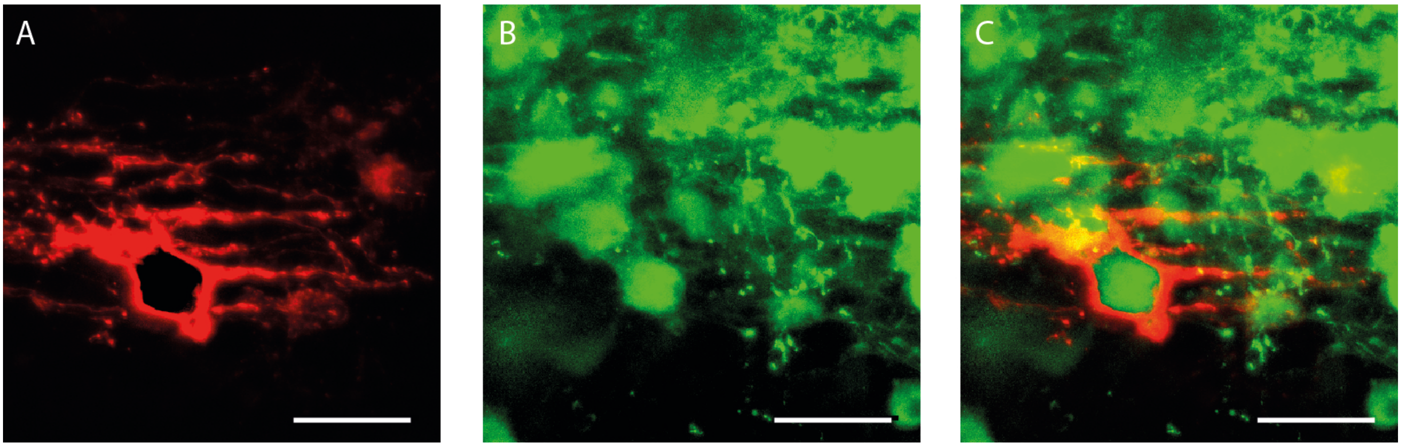

Figure S2

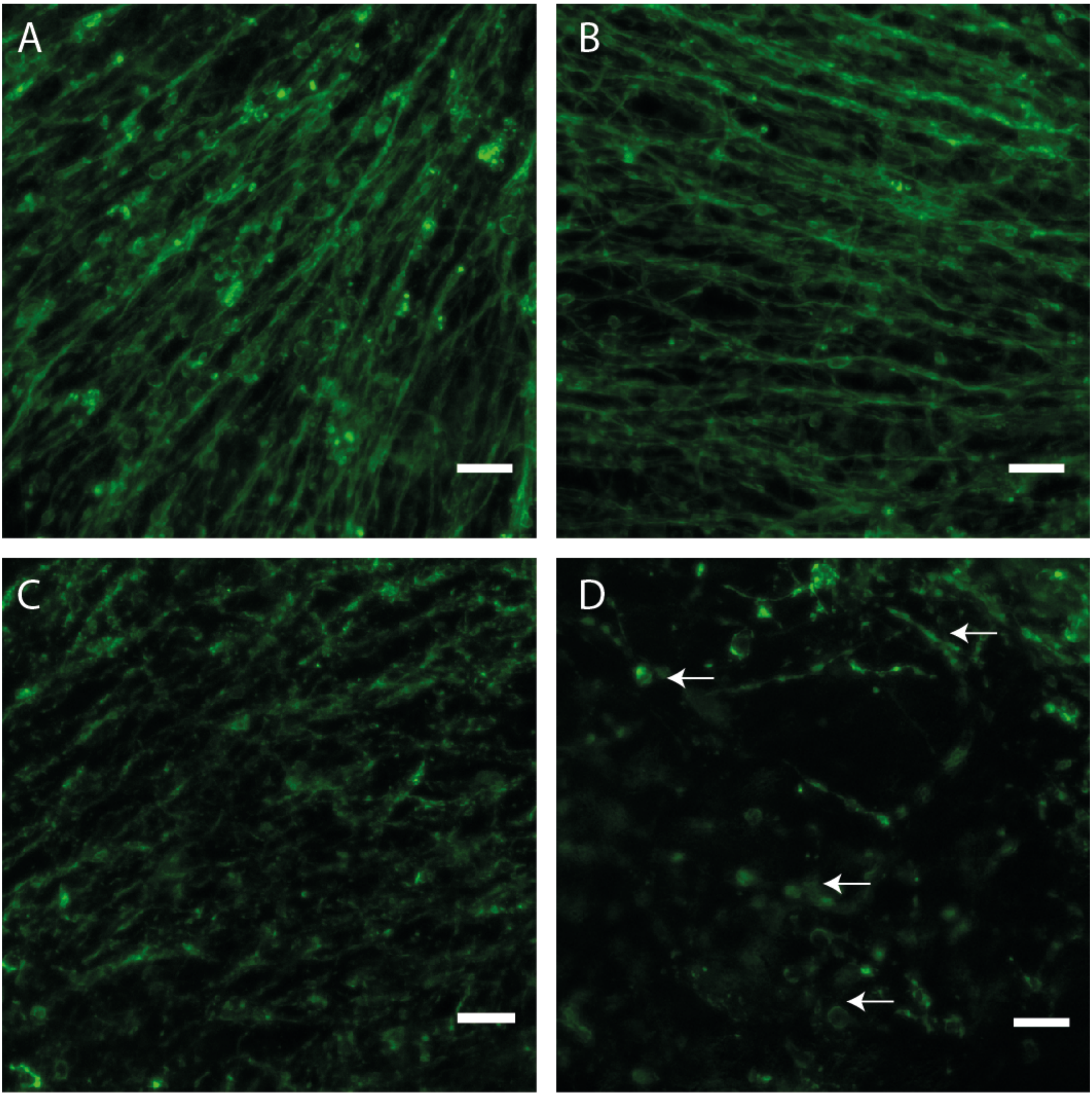

Figure S3

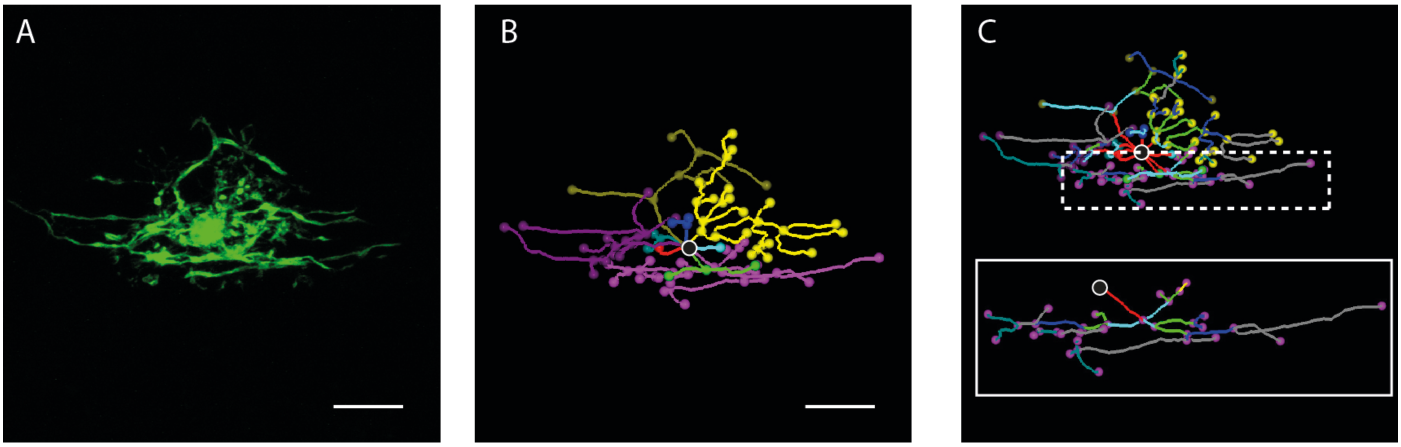

Figure S4

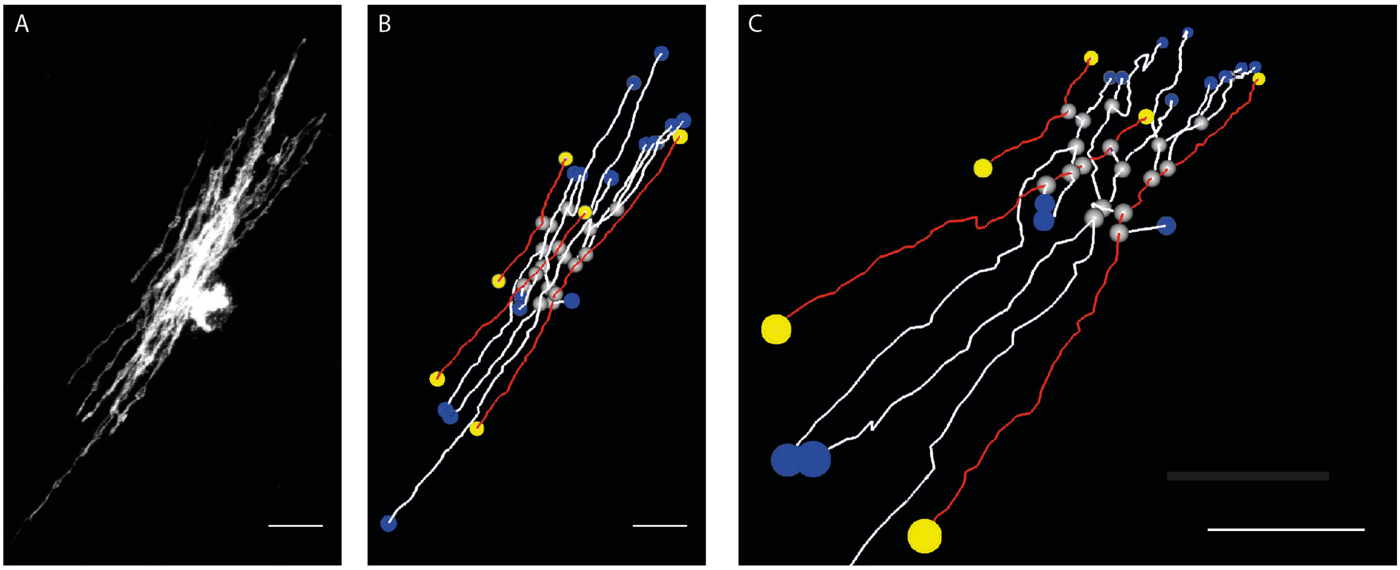
